# Epithelial cells induce the polar migration via the MHC-I–AltR interaction to eliminate transformed cells

**DOI:** 10.1101/2025.03.01.640938

**Authors:** Ryohei Teramoto, Shiyu Ayukawa, Nagisa Kamoshita, Nobuhito Goda, Takeshi Maruyama

## Abstract

In epithelial tissues, adjacent normal cells recognize and eliminate transformed cells, maintaining tissue homeostasis. We have demonstrated that the interaction between MHC-I (Major Histocompatibility Complex Class I) on transformed cells and the receptor AltR (Suboptimal Alteration Recognizing Protein) on the normal epithelial cell plays a central role in triggering the elimination ability within normal cells. Moreover, previous reports have suggested that the collective movement of surrounding normal cells including "non-adjacent peripheral normal cells" which are located further out from transformed cells also contributes to the elimination of transformed cells. However, how the direct interaction mediated by MHC-I–AltR affects the behavior of these peripheral normal cells remained unclear. In this study, we aimed to analyze the relationship between the collective movement of normal cells and the elimination of transformed cells. To achieve this, we visualized the two-dimensional movement of individual epithelial cells using the PIV (Particle Image Velocimetry) analysis and quantified the behavior of transformed cells and surrounding normal cells as vectors composed of speed and direction. As a result, we observed that a subset of normal cells near the transformed cells exhibited the unidirectional migration toward the transformed cells (polar migration), which ultimately led to their elimination. Furthermore, this polar migration was suggested to be induced by the MHC-I–AltR interaction. Additional analyses suggest that MHC-I-stimulated AltR induces Ca^2+^ signaling between normal cells, which in turn triggers the polar migration. Based on these findings, we conclude that the core molecules involved in transformed recognition, MHC-I and AltR, not only trigger the elimination ability in adjacent normal cells but also regulate the behavior of non-adjacent peripheral normal cells, contributing to the elimination of transformed cells.

## INTRODUCTION

In epithelial tissues, when transformed cells occur with a mutation in a single oncogene, surrounding normal cells eliminate them from the epithelium, thereby maintaining tissue homeostasis^1–3^. Previous studies have demonstrated that normal cells "adjacent to transformed cells" recognize MHC-I on these cells and trigger the ability to eliminate them^4^. Furthermore, recent studies have reported that the behavior of "non-adjacent surrounding normal cells," located even farther from the transformed cells, is also involved in their elimination^5–7^. However, while normal cells adjacent to transformed cells can directly recognize MHC-I via the plasma-membrane protein AltR, it remains unclear whether the behavior of the peripheral normal cells, which are located farther away, is regulated by the MHC-I–AltR interaction. To investigate whether the behavior of peripheral normal cells is regulated by the MHC-I–AltR interaction, we first developed a system to visualize the behavior of individual epithelial cells by converting their two-dimensional movement into vectors composed of speed and direction. Using this system, we analyzed how the behavior of surrounding normal cells influences the elimination of transformed cells within the epithelial monolayer. Finally, we analyzed how the interaction between MHC-I and AltR affects the behavior of surrounding normal cells.

When cells with differing states appear within the epithelium, they compete for survival. As a result, cells with higher fitness in the surrounding environment overcome as winners, while less fit cells are eliminated from the monolayer as losers. This mechanism maintains tissue homeostasis in an environment-adapted state. This phenomenon, known as cell competition, has been reported to be involved in various biological processes, including the maintenance of ES cell differentiation potential^8^, the onset of neurodegenerative diseases^9^, and the elimination of infected cells^10^. We have previously elucidated the role of cell competition in cancer prevention mechanisms in epithelial cells^4^. In a multi-step carcinogenesis model, a mutation in a single oncogene such as Ras (Rat Sarcoma Viral Oncogene) in normal epithelial cells is believed to initiate the steps of carcinogenesis. When oncogenically transformed cells appear within the epithelium, adjacent normal cells recognize these transformed cells and trigger an ability to eliminate them. Normal cells with this ability to eliminate transformed cells push them toward the apical side of the epithelium and ultimately eliminate them. In this way, the elimination of precancerous transformed cells prevents carcinogenesis.

When MHC-I on transformed cells stimulates AltR on normal cells, the normal cells trigger an ability to eliminate the transformed cells. In other words, the MHC-I–AltR interaction is one of the mechanisms underlying normal–transformed cell interactions^4^. AltR is a plasma-membrane protein that has an extracellular Ig (Immunoglobulin)-like domain for interacting with MHC-I, and an intracellular ITIM (Immunoreceptor Tyrosine-based Inhibitory Motif) domain. The human ortholog of AltR is LILRB3 (Leukocyte Immunoglobulin-Like Receptor B 3), which belongs to the LILRB family^11^. AltR on adjacent normal cells interacts with MHC-I on transformed cells. In many cancer cells, the MHC-I expression is reduced to avoid immune attacks^12^, but in early cancers or transformed cells, the MHC-I expression is enhanced compared to surrounding normal cells^4,13,14^. TCR (T-cell Receptor) recognizes the variable α1/2 domains of the MHC-I extracellular domain and peptide antigens^15^, while AltR interacts with the constant α3 domain that is recognized by NK cells and other immune cells^16^. Upon recognition of MHC-I, AltR accumulates the cytoskeletal factor Filamin at the boundary against the transformed cells via the SHP2 (Src Homology Region 2 Domain-containing Phosphatase 2)–Rho–ROCK2 (Rho-Associated Coiled-Coil Containing Protein Kinase 2) pathway, thereby pushing the transformed cells toward the apical side. In this manner, adjacent normal cells, through the MHC-I–AltR interaction, accumulate Filamin as a force to eliminate the transformed cells. Furthermore, in addition to the accumulation of Filamin, it has been reported that peripheral normal cells, located outside the adjacent cells, collectively move toward the transformed cells, contributing to their elimination^5^. However, the molecular mechanisms regulating this collective movement and whether the MHC-I–AltR interaction regulates the collective behavior of surrounding normal cells remain unclear.

The elimination of cells from the epithelium is triggered by cell-cell interactions. These interactions are categorized into physical interactions resulting from cell contact and chemical interactions such as ligand-receptor interactions^17^. Previous reports have shown that physical distortions arising within the cell layer, due to processes like cell proliferation, induce the movement of specific cell populations toward particular cells. This leads to the elimination of suboptimal cells from the monolayer. In other words, the cell elimination resulting from physical interactions is defined as "passive elimination." On the other hand, chemical interactions are the core mechanism behind cell competition, and cell elimination resulting from chemical interactions is defined as "active elimination." A key point in the existing consensus is that passive elimination involves distortions within the monolayer and leads to the movement of surrounding cells, whereas in active elimination, only the cells adjacent to the eliminated cells acquire the ability to eliminate them. Interestingly, it has recently been reported that when arise, normal cells surrounding Src-transformed cells induce the collective movement^18^. This suggests that, not only in passive elimination but also in active elimination surrounding normal cells may undergo collective behavioral changes^5,19^. Additionally, in the case of Ras-transformed cell elimination, it has been suggested that the Ca^2+^ signal originating from Ras-transformed cells induces collective behavioral changes in surrounding normal cells, supporting the hypothesis that collective behavioral changes in surrounding normal cells, not just those adjacent to the transformed cells, contribute to active elimination^6^.

In the phenomenon of cell competition, it has been suggested that the transduction of the Ca^2+^ signal originating from transformed cells triggers the collective movement of surrounding epithelial cells toward the transformed cells^6^. In transformed cells, the TRPC (Transient Receptor Potential Channel), a Ca^2+^ channel in the cell membrane, takes up extracellular Ca^2+^, while the IP_3_R (Inositol 1,4,5-Trisphosphate Receptor), a Ca^2+^ channel in the endoplasmic reticulum (ER), releases Ca^2+^ from the ER. The increased Ca^2+^ in the transformed cells flows into adjacent normal cells through gap junctions between transformed and normal cells. In response to this influx of Ca^2+^, adjacent normal cells also increase intracellular Ca^2+^ via a similar response to the transformed cells. Ca^2+^ is then transduced to surrounding normal cells through gap junctions between normal cells, even those not adjacent to the transformed cells. Furthermore, actin rearrangement occurs within these normal cells, leading to a change in their direction toward the transformed cells. These observations suggest that the Ca^2+^ signal originating from transformed cells plays an important role in cell-cell interactions, triggering the collective movement of surrounding normal cells. However, how this Ca^2+^ signal from transformed cells is regulated, and whether chemical interactions, such as the MHC-I–AltR interaction that plays a crucial role in active elimination, influence this process, has not been analyzed in detail.

In this study, we show that the collective movement of surrounding normal cells specifically triggered by transformed cells leads to the elimination of transformed cells from the epithelium. Additionally, we demonstrate that the MHC-I–AltR interaction mediates the Ca^2+^ signal and induces collective behavioral changes.

## RESULTS

### Surrounding Normal Cells Eliminate Transformed Cells via the Polar Migration

To examine whether surrounding normal cells exhibit the collective movement toward transformed cells, we co-cultured normal human skin epithelial cells, HaCaT cells, with HaCaT-pTRE3G-GFP-RasV12 (HaCaT-RasV12) cells expressing doxycycline-inducible GFP-RasV12 and observed the cell layer using live imaging with both bright-field and fluorescence microscopy (**Figure 1A**). The behavior of individual transformed cells and their surrounding normal cells was visualized by converting the movement speed and direction into vectors using PIV analysis. Cells with larger mobility were indicated with red arrows, while those with smaller mobility were indicated with blue arrows. As a result, 20 hours after doxycycline stimulation, cells that were slowly moving, as indicated by the blue arrows, came to a near pause at 24 hours. However, by 28 hours, normal cells surrounding RasV12 cells collectively moved significantly toward RasV12 cells, as indicated by the red arrows, suggesting that this collective movement was unidirectional, “polar migration.” At 32 hours, RasV12 cells were eliminated, and by 34 hours, when the RasV12 cells were completely eliminated from the epithelial monolayer, the group of normal cells moving toward RasV12 cells disappeared (**Figure 1B**). This suggests that, in the epithelial monolayer, surrounding normal cells exhibit the polar migration to eliminate RasV12 cells. Furthermore, to investigate whether there is a correlation between the polar migration and the elimination of RasV12 cells, we performed regression analysis with the number of surrounding normal cells exhibiting the polar migration between 20-50 hours on the x-axis and the elimination efficiency of RasV12 cells on the y-axis. The results showed that both the number of surrounding normal cells exhibiting the polar migration and the elimination efficiency of RasV12 cells increased over time. The R^2^ value indicating correlation was 0.97 (**Figure 1C**). These results suggest that surrounding normal cells induce the polar migration to eliminate transformed cells.

**Figure 1.**
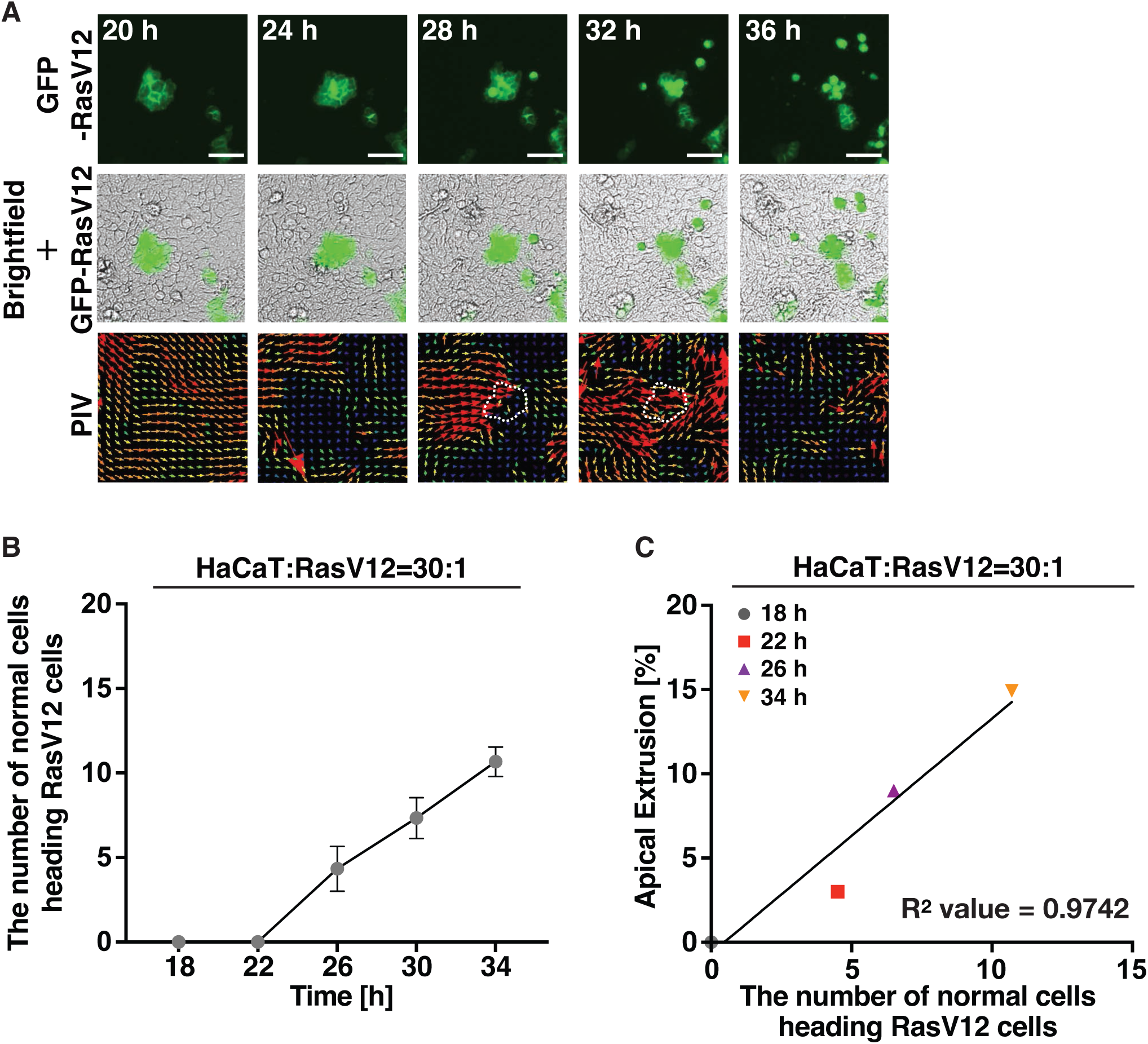
Surrounding Normal Cells Eliminate Transformed Cells via the Polar Migration. (**A**) Imaging of the epithelial monolayers during co-culture. HaCaT cells and HaCaT-RasV12 cells were mixed in a 30:1 ratio and seeded onto a thin collagen layer to form a monolayer. GFP-RasV12 expression was induced by doxycycline stimulation, and live imaging was conducted every 5 minutes from 8 hours after induction for 48 hours. The upper panel shows brightfield images captured using a light microscope, the middle panel shows GFP fluorescence images, and the lower panel shows PIV images. The time notation indicates the elapsed time after GFP-RasV12 expression induction. The dashed line indicates the position of HaCaT-RasV12 cells at 28 hours. PIV analysis was performed on brightfield images obtained at 2 hours using the ImageJ plugin. Red and purple arrows indicate high and low mobility, respectively. Scale bars: 100 μm. (**B**) MHC-I-AltR interaction induces polar migration. A graph showing the number of normal cells exhibiting polar migration, as indicated by the red arrows. Data are mean ± SEM. n > 375 cells. (**C**) There was a strong positive correlation between polar migration and the elimination of transformed cells. The R^2^ value, calculated from regression analysis, was compared with the threshold value of 0.7, indicating a strong correlation. n > 375 cells.

### The MHC-I–AltR Interaction Eliminates Transformed Cells through the Polar Migration

Next, we investigated whether the polar migration is regulated by the molecular interaction between MHC-I and AltR, which are involved in the recognition of transformed cells. We previously demonstrated that the interaction between MHC-I on RasV12 cells and AltR on normal cells leads to the elimination of RasV12 cells and that treatment with recombinant protein of MHC-I-α3 (rec.α3), an interacting domain of MHC-I, promotes the elimination of RasV12 cells, while knockout (KO) of AltR suppresses the elimination of RasV12 cells regardless of the presence of rec.α3^4^. To further investigate this, we observed the behavior of surrounding normal cells using rec.α3 and normal HaCaT-LILRB3 KO cells, which lack the human ortholog gene of AltR. The results showed that, in the absence of rec.α3, the polar migration of normal cells was unidirectional, whereas the rec.α3 treatment resulted in a shift to the multidirectional polar migration, with an increased number of normal cells migrating toward RasV12 cells (**Figure 2A, B**). To quantify the number of normal cells showing the polar migration, we measured the cells exhibiting the polar movement toward RasV12 cells, indicated by red arrows, and displayed this in a polar coordinate histogram centered on RasV12 cells. As a result, unidirectional polar migration in the untreated condition shifted to multiple directions with the rec.α3 treatment, showing a higher number of polar migration events from various directions (**Figure 2C**). On the other hand, LILIRB3 KO in normal cells suppressed both the polar migration and the elimination of RasV12 cells regardless of the rec.α3 treatment (**Figure 2A-C**). These findings suggest that the MHC-I–AltR interaction induces the polar migration. Quantification of polar migration revealed that, at 34 hours, around 10 surrounding normal cells exhibited polar migration in HaCaT-wild-type (WT) cells, while the rec.α3 treatment increased this to approximately 15 cells. However, LILRB3 KO in normal cells reduced the number of cells showing polar migration to about 5, and the rec.α3 treatment did not enhance migration (**Figure 2D**). Regarding the elimination efficiency of RasV12 cells, HaCaT-WT cells exhibited a 30% elimination rate, which increased to about 40% with the rec.α3 treatment. In contrast, LILRB3 KO in normal cells suppressed the elimination efficiency to approximately 5%, and the rec.α3 treatment did not promote RasV12 cell elimination (**Figure 2E**). Furthermore, regression analysis of the number of normal cells showing the polar migration and the elimination efficiency of RasV12 cells revealed a strong positive correlation, with an R^2^ value of 0.90, which is significantly higher than 0.7 (**Figure 2F**). These results suggest that the MHC-I–AltR interaction eliminates transformed cells through the polar migration.

**Figure 2.**
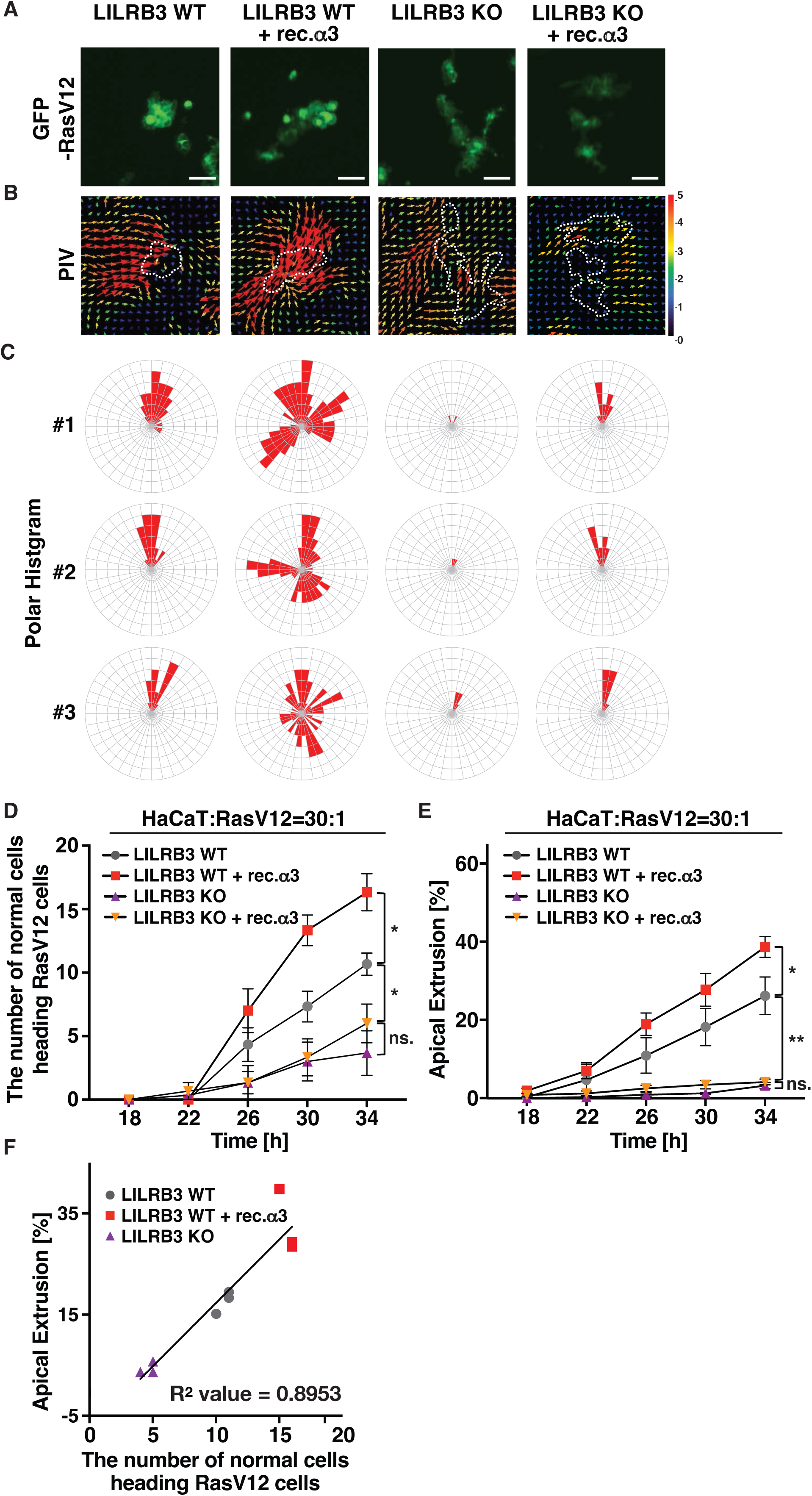
The MHC-I–AltR Interaction Eliminates Transformed Cells through the Polar Migration. (**A and B**) MHC-I–AltR interaction induces polar migration. Fluorescence images of GFP (**A**) and the corresponding PIV images (**B**) are shown. HaCaT-wild type (WT) or - LILRB3 knockout (KO) cells were mixed with HaCaT-RasV12 cells in a 30:1 ratio and seeded onto a thin collagen layer. After monolayer formation, GFP-RasV12 expression was induced by doxycycline stimulation, and recombinant MHC-I α3 protein (rec.α3) was added. Live imaging was conducted every 5 minutes from 8 hours after induction for 48 hours. The time notation indicates the elapsed time after GFP-RasV12 induction. The dashed line indicates the position of RasV12 cells at 28 hours. PIV analysis was performed on brightfield images obtained at 2 hours using the ImageJ plugin. Red and purple arrows indicate high and low mobility, respectively. Scale bars: 100 μm. (**C**) Polar coordinate histogram based on PIV images. Red arrows showing polar migration toward HaCaT-RasV12 cells were measured, with the part where the most red arrows appeared set as 0 degrees. (**D and E**) MHC-I–AltR interaction induces both polar migration and elimination of transformed cells. Graphs showing the number of normal cells exhibiting polar migration (**D**) and the elimination efficiency (**E**). Data are mean ± SEM. n > 375 cells. \**P* < 0.05, \*\**P* < 0.01 by one-way ANOVA. ns.: not significant. (**F**) There was a strong positive correlation between the polar migration and the elimination of transformed cells. The R^2^ value, calculated from regression analysis, was compared with the threshold value of 0.7, indicating a strong correlation. n > 375 cells.

### The Polar Migration is Induced by the Ca^2+^ Signals to Surrounding Normal Cells

Next, we investigated how surrounding normal cells interact with each other to induce the polar migration. It has been reported that MHC-I-stimulated AltR accumulates the skeletal formation factor Filamin at the boundary against RasV12 cells^4^. We examined whether the Filamin accumulation via the MHC-I–AltR interaction is involved in the induction of the polar migration. As a result, Filamin knockdown (KD) in normal cells did not change the number of normal cells exhibiting the polar migration (**Figure 3A-D**). However, the elimination efficiency of RasV12 cells was suppressed by Filamin KD (**Figure 3E**). These results suggest that the Filamin accumulation via the MHC-I–AltR interaction is necessary for the elimination of RasV12 cells but not involved in the induction of polar migration. In other words, it is suggested that surrounding normal cells may induce the polar migration by transducing some form of signal.

**Figure 3.**
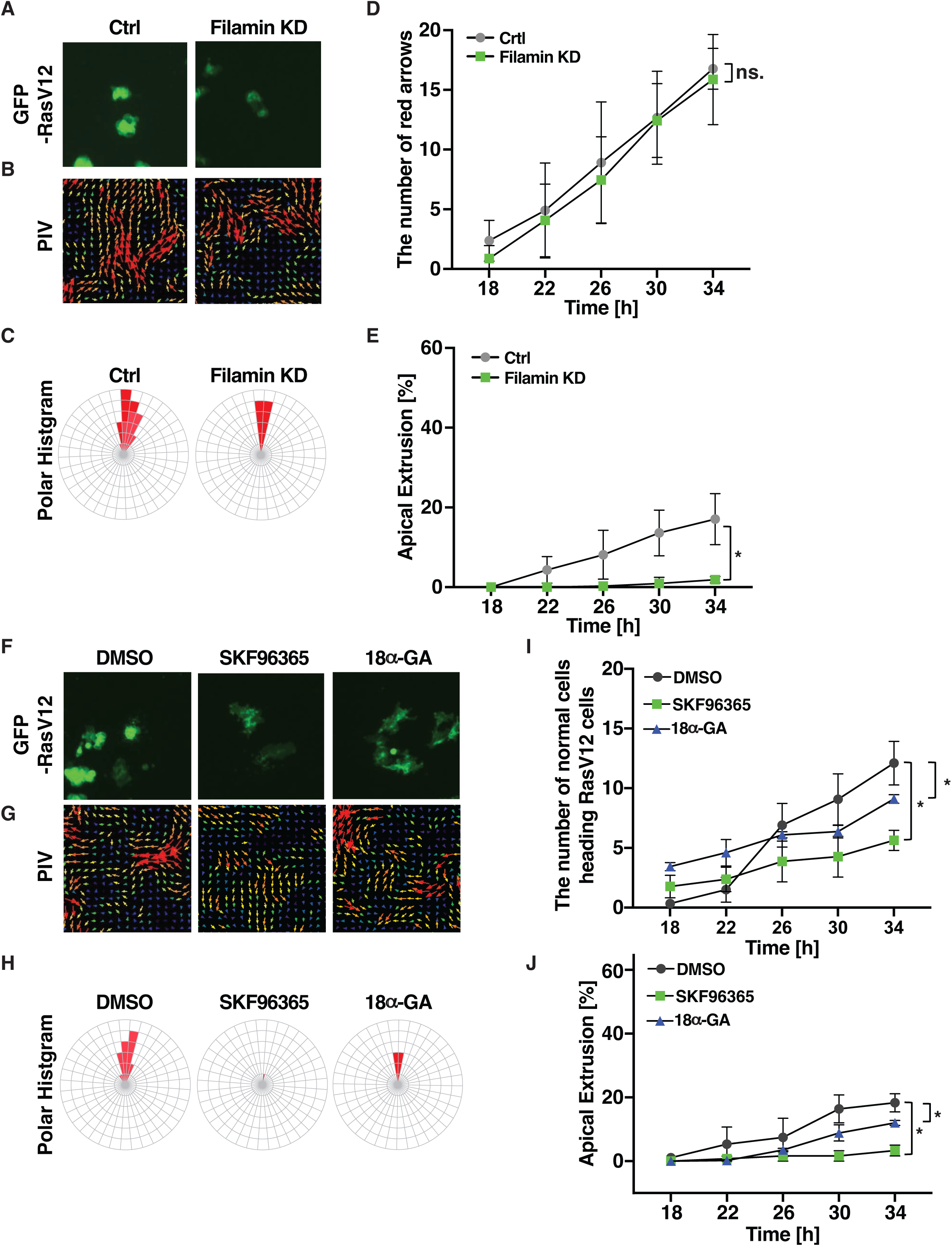
The Polar Migration is Induced by the Ca^2+^ Signals to Surrounding Normal Cells. (**A, B, F and G**) Fluorescence images of GFP (**A, F**) and the corresponding PIV images (**B, G**) are shown. HaCaT-WT cells were mixed with RasV12 cells or RasV12-Filamin knockdown (KD) cells (**A, B**) in a 30:1 ratio and seeded onto a thin collagen layer. After monolayer formation, GFP-RasV12 expression was induced by doxycycline stimulation, and each inhibitor was added (**F, G**). Time notation indicates the elapsed time after GFP-RasV12 induction. PIV analysis was performed on brightfield images obtained at 2 hours using the ImageJ plugin. Red and purple arrows indicate high and low mobility, respectively. The inhibitors were added at the following concentrations: SKF96365: 50 mM; 18 alpha-GA: 100 μM. (**C and H**) Polar coordinate histogram based on PIV images. Red arrows showing polar migration toward HaCaT-RasV12 cells were measured. (**D, E, I and J**) Graphs showing the number of normal cells exhibiting polar migration (**D, I**) and the elimination efficiency (**E, J**). Data are mean ± SEM. n > 280 cells. \**P* < 0.05 by Student’s *t*-tests. ns.: not significant.

Previous studies have suggested that Ca^2+^ waves generated in RasV12 cells propagate to surrounding normal cells, leading to their collective migration toward RasV12 cells^6^. To elucidate the signals transduced by surrounding normal cells that induce the polar migration, we focused on the behavioral changes of surrounding normal cells induced by Ca^2+^ signals originating from RasV12 cells. To test whether Ca^2+^ signals regulate the polar migration and the elimination of RasV12 cells, we inhibited molecules responsible for the influx of extracellular Ca^2+^, such as TRPC, and molecules responsible for the influx of Ca^2+^ from adjacent cells via Gap Junctions. The results showed that treatment with TRPC inhibitors and Gap Junction inhibitors reduced the number of normal cells migrating toward RasV12 cells (**Figure 3F-I**). In other words, both TRPC inhibitors and Gap Junction inhibitors suppressed the polar migration. Furthermore, the elimination efficiency of RasV12 cells was also suppressed by treatment with either TRPC inhibitors or Gap Junction inhibitors (**Figure 3J**). These findings suggest that Ca^2+^ signals induce the polar migration, which is necessary for the elimination of transformed cells.

### The MHC-I–AltR Interaction Induces the Polar Migration by Transducing Ca^2+^ Signals to Surrounding Normal Cells

To investigate whether Ca^2+^ signals from RasV12 cells are transduced to surrounding normal cells, we performed imaging of Ca^2+^ signals using Rhod-5N, a Ca^2+^ indicator dye. As a result, 18 hours after RasV12 expression induction, the fluorescence of Rhod-5N was observed within RasV12 cells. By 22 hours, the fluorescence of Rhod-5N was observed in two adjacent normal cells. These results suggest that the initially activated Ca^2+^ signal in RasV12 cells was transduced to the adjacent normal cells (**Figure 4A**). Next, we tested whether this transduction of Ca^2+^ signals was regulated by the MHC-I– AltR interaction, using rec.α3 and AltR KO normal cells. The results showed that after the rec.α3 treatment, the Ca^2+^ signal generated in RasV12 cells was transduced to a higher number of more distant surrounding normal cells compared to untreated conditions. Additionally, when AltR was knocked out in normal cells, the Ca^2+^ signal was still generated in RasV12 cells, but it was only transduced to adjacent normal cells, regardless of the rec.α3 treatment (**Figure 4B**). To quantitatively analyze the Ca^2+^ signal, we measured the fluorescence intensity of Rhod-5N based on images obtained by imaging. The fluorescence intensity of Rhod-5N in WT normal cells increased statistically significantly after the rec.α3 treatment, while AltR KO in normal cells significantly decreased. Furthermore, when rec.α3 was treated to AltR KO normal cells, the fluorescence intensity was significantly lower compared to WT normal cells treated with rec.α3 (**Figure 4C**). These findings suggest that the MHC-I–AltR interaction positively regulates the transduction of Ca^2+^ signals among normal cells. Taken together, these results suggest that the MHC-I–AltR interaction induces the polar migration and eliminates transformed cells via Ca^2+^ signal transduction.

**Figure 4.**
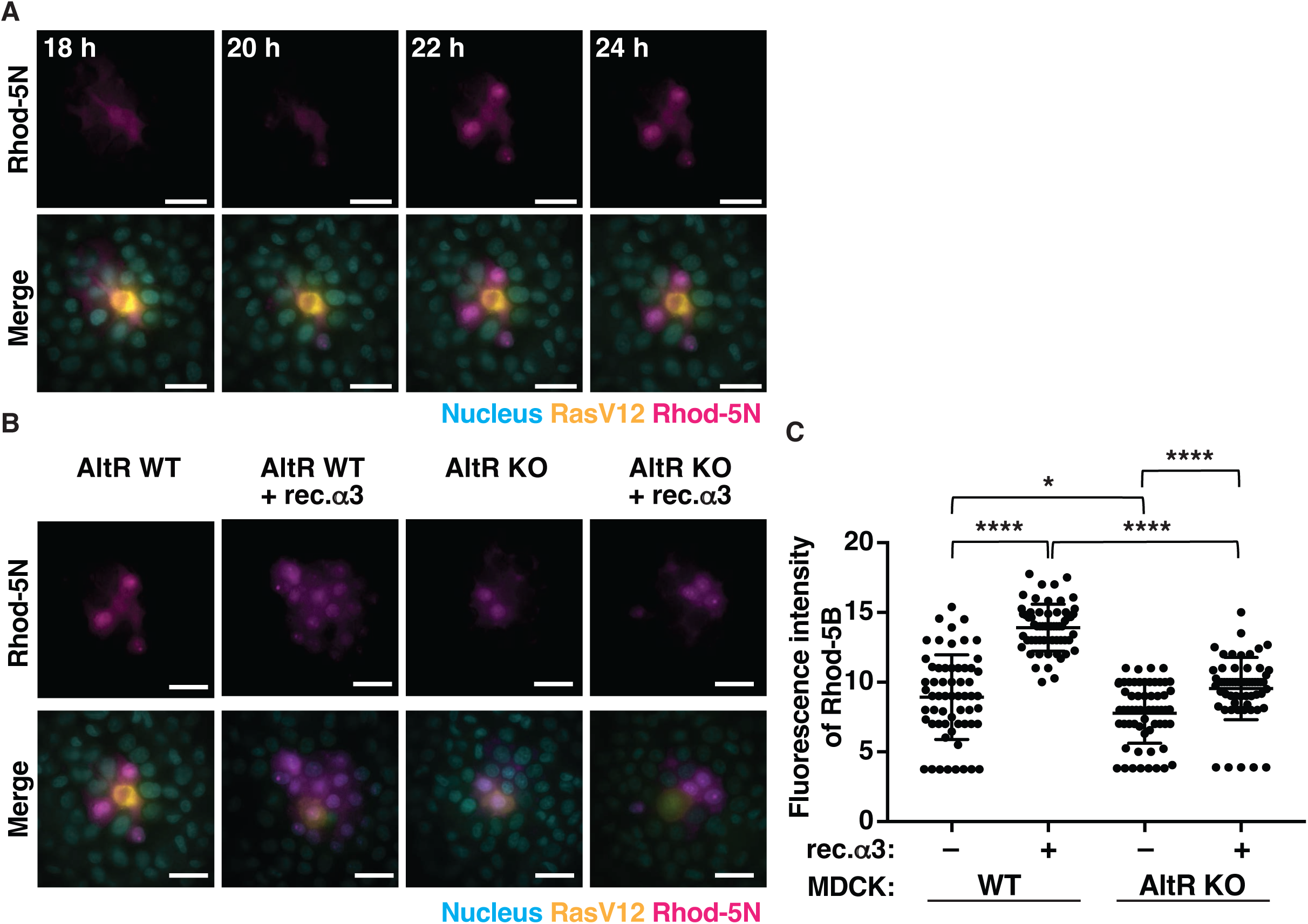
The MHC-I–AltR Interaction Induces the Polar Migration by Transducing Ca^2+^ Signals to Surrounding Normal Cells. (**A and B**) The MHC-I–AltR interaction induces Ca^2+^ signals. MDCK-WT or -AltR KO cells (**B**) were mixed with MDCK-RasV12 cells in a 30:1 ratio and seeded onto a thin collagen layer. After the monolayer formation, GFP-RasV12 expression was induced, and rec.α3 was added (**B**). After 16 hours of culture, nuclei were stained with Hoechst33342 (blue). Rhod-5N (magenta) was added, and live imaging was performed every 3 minutes for 18 hours using a disc-scanning microscope. The upper image shows the fluorescence image of Rhod-5N, and the lower image shows the overlay of Rhod-5N, GFP-RasV12, and nuclei. Time notation indicates the elapsed time after GFP-RasV12 induction. Scale bars: 20 μm. (**C**) Graph showing the signal intensity of Rhod-5N in peripheral normal cells. In the images (B), a circle with a radius of 5 cells was drawn from the MDCK-RasV12 cells, and the area inside was used as the measurement region. Data are mean ± SD. n > 180 cells. \**P* < 0.05, \*\*\*\**P* < 0.001 by Student’s *t*-tests. ns.: not significant.

## DISCUSSION

In this study, we demonstrated that MHC-I and AltR, which were previously revealed as key players in the interaction between normal and transformed cells, induce the polar migration of surrounding normal cells that are not adjacent to the transformed cells. Furthermore, we observed that polar migration occurs through the transduction of the Ca^2+^ signal in the sequence of transformed cells → adjacent normal cells → peripheral normal cells, leading to the elimination of transformed cells. In this transduction of the Ca^2+^ signal, MHC-I-stimulated AltR seems to regulate the Ca^2+^ signal between adjacent and peripheral normal cells. These findings suggest that the MHC-I–AltR interaction regulates not only the behavior of adjacent normal cells but also that of surrounding normal cells that are not in direct contact with the transformed cells, thereby eliminating the transformed cells. This represents a novel mechanism of transformed cell elimination mediated by the MHC-I–AltR interaction. In addition, the Filamin suppression in normal cells did not affect the polar migration, but it inhibited the elimination of transformed cells. Therefore, the MHC-I–AltR interaction is thought to induce the polar migration in surrounding normal cells, and the stress generated by this migration is transduced to transformed cells via Filamin accumulated in adjacent normal cells, leading to their elimination.

This study suggests that AltR induces the Ca^2+^ signal in surrounding normal cells. However, the mechanisms regulating the influx and efflux of Ca^2+^ by AltR in normal cells remain unclear. When AltR was deleted in normal cells, the Ca^2+^ signal from transformed cells was only transduced to adjacent normal cells. This suggests that AltR may regulate the influx and efflux of Ca^2+^ within adjacent normal cells and the propagation of Ca^2+^ signals between adjacent and peripheral normal cells. Additionally, since both TRPC and Gap Junction inhibitors suppressed the elimination of transformed cells and the polar migration of normal cells, these molecules and mechanisms are likely involved downstream of AltR. Furthermore, SHP2 is recruited to the phosphorylated IL-1 receptor, and activated SHP2 binds to IP_3_R, suppressing the release of Ca^2+^ from the ER^20^. Future studies should investigate whether AltR is involved in the activation of IP_3_R and other Ca^2+^ regulatory factors.

In the *in vitro* cell competition model, the polar migration is induced by the MHC-I–AltR interaction. However, this polar migration was not observed in all adjacent normal cells but only in a subset of adjacent normal cells. This suggests that normal cells lacking the MHC-I–AltR interaction signal do not undergo polar migration. One possible reason for this could be differences in the expression levels of AltR in adjacent normal cells. For example, if the expression level of AltR is insufficient to interact with MHC-I, the polar migration may not occur. In this study, the rec.α3 treatment led to more adjacent normal cells showing the polar migration. This suggests that rec.α3 can provide the MHC-I signal even to adjacent normal cells that express AltR but not at sufficient levels to respond to MHC-I in RasV12 cells.

Moreover, the expression of AltR in adjacent normal cells is induced by Runx2 activated via the detection the increased membrane tension of neighboring transformed cells^4^. It is presumed that the intensity of the Runx2 signal determines the difference in the expression levels of AltR. Therefore, we anticipate that differences in the contact area between normal cells and transformed cells will result in variations in AltR expression levels. Future studies will focus on the correlation between AltR expression levels and polar migration, considering the contact area between transformed cells and normal cells.

This study shows that normal cells recognizing MHC-I on transformed cells induce the polar migration through interactions between normal cells via AltR, leading to the elimination of transformed cells. Therefore, in mammalian cell competition, the elimination of transformed cells requires not only the interaction between transformed cells and adjacent normal cells but also the interactions between surrounding normal cells.

## MATERIALS AND METHODS

### Cell Culture

Madin-Darby Canine Kidney (MDCK), MDCK-AltR KO, MDCK pTRE3G-GFP-RasV12 (MDCK-RasV12), HaCaT, HaCaT-LILRB3 KO and HaCaT pTRE3G-GFP-RasV12 (HaCaT-RasV12) cells were cultured as previously described^4^. To induce the expression of GFP-RasV12, the doxycycline-inducible MDCK- and HaCAT-RasV12 cell lines were treated with 100 ng/mL doxycycline (Sigma-Aldrich) and cultured for 16 (MDCK-RasV12 cells) or 8 (HaCaT-RasV12 cells) hours.

### Reagents

A gap junction inhibitor, 18 alpha-glycyrrhetinic acid (18α-GA), and DMSO were purchased from Sigma-Aldrich. A TRPC inhibitor, SKF96365, was from Funakoshi. A Ca^2+^ indicator, Rhod-5N, was from Thermo Fisher Scientific. Each inhibitor was used at the indicated concentration below: SKF96365 50 mM for HaCaT cells; 18α-GA 100 μM for HaCaT cells; Rhod-5N 30 μM for MDCK cells.

### Recombinant protein purification

pGEX-6P-α3-Flag^4^ was transformed into BL21 competent cells, which were then plated onto LB agar plates supplemented with ampicillin. After incubation at 37°C for 16 hours, individual colonies were picked and cultured in LB medium containing ampicillin at 37°C for an additional 16 hours. A portion of the culture was transferred to 50 mL of fresh LB medium with ampicillin (20 ng/mL) and incubated at 37°C for 16 hours. The culture was subsequently scaled up by transferring to 450 mL of ampicillin-containing LB medium (20 ng/mL). Once the OD600 of the culture reached 0.5, the pre-cultured cells were induced with 0.1 mM isopropyl β-D-thiogalactopyranoside (IPTG) and cultured at 18°C for 24 hours. Cells were harvested by centrifugation, resuspended in 1% Triton X-100 lysis buffer (10 mM EDTA/PBS) supplemented with Aprotinin, and subjected to sonication (5 cycles of 45% power for 1 minute using a UD-100 sonicator, TOMY). Finally, the cell lysates were purified using glutathione-conjugated Sepharose 4B beads (GE Healthcare), and the recombinant proteins were eluted with 10 mM reduced glutathione buffer (50 mM Tris-HCl, pH 8.0).

### Live imaging for *in vitro* cell competition model system

HaCaT-RasV12 cells were mixed with normal HaCaT-WT or -LILRB3 KO cells at a ratio of 1:30 and cultured on the thin collagen matrix as previously described^4^. The mixed cells were cultured for 16 hours until the formation of a monolayer, followed by doxycycline treatment for the indicated time. SKF96365, 18α-GA or rec.α3 was added with doxycycline at the indicated concentration. Live imaging was conducted every 5 minutes from 8 hours after the doxycycline treatment for 48 hours with JuLI Stage (NanoEntek).

The cell elimination efficiency was calculated based on the bright-field and fluorescence images obtained from live imaging. The imaging data acquired over 2 hours were converted into a video using ImageJ, and the cell behavior was observed over time. Cells that became round and moved randomly with significant displacement were considered eliminated cells. The extrusion observation period was defined as the 30-hour interval from 20 hours after the doxycycline treatment, when the movement of the cell layer paused, to 50 hours, when the elimination of normal cells was first observed. Additionally, the total number of cells in the field of view was measured using the CellProfiler, and the ratio of eliminated cells to the total number of cells in the field of view was calculated as the elimination efficiency.

### Short interference RNA transfection into normal cells

HaCaT cells were transiently transfected with RNAi (30 pmol) using Lipofectamine RNAiMAX (Invitrogen), following the manufacturer’s instructions. The Stealth RNAi siRNA Negative Control, Med GC (Invitrogen), was used as a negative control. After 48 hours, the siRNA-transfected cells were mixed with HaCaT-RasV12 cells and seeded onto thin collagen gels. The sampling procedure is detailed in “Live imaging for *in vitro* cell competition model system.” The sequences of RNAi are as follows:

siFilamin#1: 5’-GAUGGCGUGUAUGGCUUCGAGUAUU-3’; siFilamin#2: 5’-GAGGCCAAGAUGUCCUGCAUGGAUA-3’.

### Particle Image Velocimetry (PIV) analysis

To perform PIV analysis, 24 bright-field images obtained over 2 hours through live imaging were compiled into a video using ImageJ’s image sequence function. Based on this video, the PIV plugin was used to display the individual cell movement speed and direction with arrows. Red arrows indicate high mobility, while purple arrows indicate low mobility. The parameters for the PIV analysis are as follows: Normalized median test parameters noise for NMT (0.3); threshold for NMT (0.8); Dynamics mean test parameters C1 for DMT (0.5); C2 for DMT (0.3).

### Creation of Polar Coordinate Histogram

To quantify the number of normal cells showing the polar migration, the PIV analysis results were converted into a polar coordinate histogram centered around the transformed cells. The transformed cells at the apex of the polar migration were placed at the center of the polar coordinates, with the coordinates divided into 10-degree intervals. The number of red arrows, representing normal cells showing the polar migration, was counted within each sector and displayed as red markers. The polar coordinates were rotated such that the sector with the highest number of red arrows was set to 0 degrees.

### Live Imaging of Ca^2+^ Signals

MDCK-WT or -AltR KO cells were mixed with MDCK-RasV12 cells at a 30:1 ratio, seeded onto a thin collagen layer, and cultured at 37°C for 24 hours to form a monolayer. After induction of GFP-RasV12 expression with doxycycline, the cells were cultured for an additional 16 hours. Hoechst 33342 was diluted 1:1000 and used to stain the nuclei for 10 minutes. After adding Rhod-5N to the culture medium, imaging was performed every 3 minutes for 18 hours using a disc-scanning microscope (Olympus) at 42°C with 5% CO_2_.

Fluorescence intensity of Rhod-5N was measured based on the fluorescence images obtained from disc-scanning microscopy. The measurement was performed using ImageJ. In the group with the rec.α3 treatment, normal cells located within a distance of approximately 5 cells from the transformed cells exhibited the polar migration. Therefore, in the Ca^2+^ signal live imaging, a circle with a radius corresponding to a distance of 5 cells from the transformed cells was drawn, and the area inside the circle was set as the region of measurement. The fluorescence intensity of Rhod-5N was measured at 66 points within this region.

### Quantification and Statistical Analysis

Statistical analyses were performed using Prism 7.0, and p-values were determined by two-tailed Student’s *t*-test or one-way ANOVA. Data obtained from three or more independent experiments are presented as the mean ± standard error of the mean (mean ± SEM) or mean ± standard deviation (mean ± SD). The elimination efficiency of transformed cells in live imaging was calculated based on a sample of more than 375 transformed cells. For the quantification of Rhod-5N fluorescence intensity, fluorescence intensities in transformed cells and surrounding normal cells were analyzed using ImageJ for more than 50 cells. Additionally, to demonstrate correlation, regression analysis was performed using the statistical analysis tool R, and the R² value was calculated.

## AUTHOR CONTRIBUTIONS

Conceptualization, T.M.; Investigation, R.T. and N.K.; Writing – Original Draft, R.T., S.A. and T.M.; Writing – Review & Editing, S.A. and T.M.; Funding Acquisition, T.M.; Supervision, N.G. and T.M.

## ACKNOWLEDGMENT

This work was supported by Japan Society for the Promotion of Science (JSPS) Grant-in-Aid for Scientific Research (B) (Grant Number 18H02675), the Fusion Oriented REsearch for disruptive Science and Technology (FOREST) (Grant Number JPMJFR226D) from the Japan Science and Technology Agency (JST), Advanced Research & Development Programs for Medical Innovation (Prime) (Grant Number 19gm6210019h0001) and Project for Promotion of Cancer Research and Therapeutic Evolution (P-PROMOTE) (Grant Number 23ama221113h0002) from the Japan Agency for Medical Research and Development (AMED), Takeda Science Foundation, the Naito Foundation and the TERUMO Life Science foundation (to TM).

